## Supplementary material for "Dimeric ^R25C^PTH(1-34) Activates the Parathyroid Hormone-1 Receptor *in vitro* and Stimulates Bone Formation in Osteoporotic Female Mice": Figure supplement 1

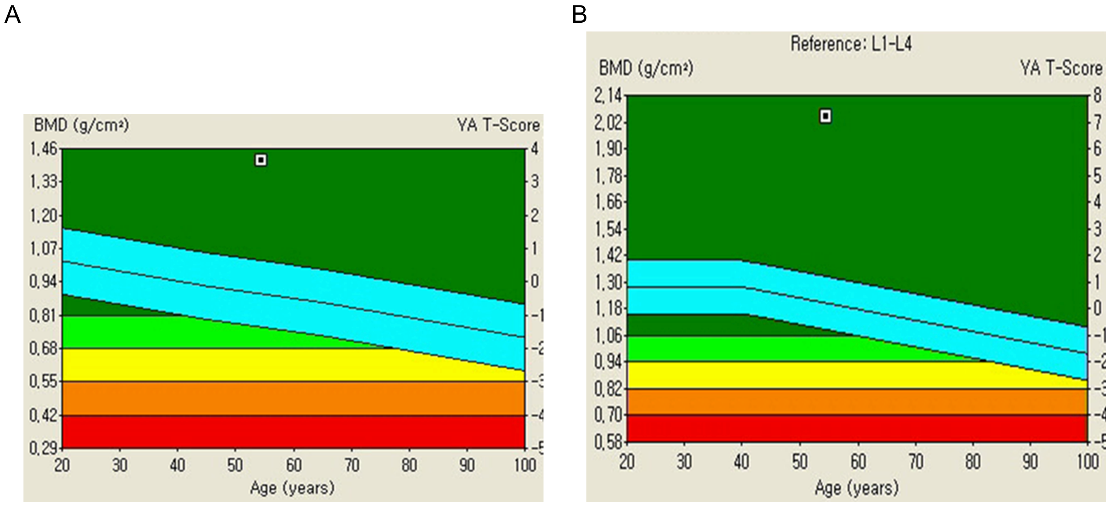

**Figure supplement 1.** Bone Mineral Density (BMD) Analysis in a Patient Affected by Homogeneous ^R25C^PTH Mutation.

Trabecular bone score (TBS) is a measure in bone density assessments, evaluating bone tissue microarchitecture. Derived from dual-energy X-ray absorptiometry (DXA) scans, TBS aids in fracture risk determination. The T-score, present on bone density reports, reflects bone mass deviation from the norm within the same age group. (A) The TBS report demonstrating a significantly elevated BMD in the lumbar spine of the patient. (B) The TBS report demonstrating a significantly elevated BMD in the femur of the patient. The left Y-axis indicates BMD (g/cm2), while the right Y-axis presents T-score. T-score interpretation is as follows. Normal or lower risk of fractures: -1.0 and above; Low bone density or osteopenia: Between -1.0 and -2.5, indicating weaker bones than normal but not classified as osteoporotic; Osteoporosis: -2.5 and below, signifying a substantial fracture risk due to very low bone density. Light blue indicates average T-score of normal population ± 1.
