## Supplementary material for "Dimeric ^R25C^PTH(1-34) Activates the Parathyroid Hormone-1 Receptor *in vitro* and Stimulates Bone Formation in Osteoporotic Female Mice": Figure supplement 2

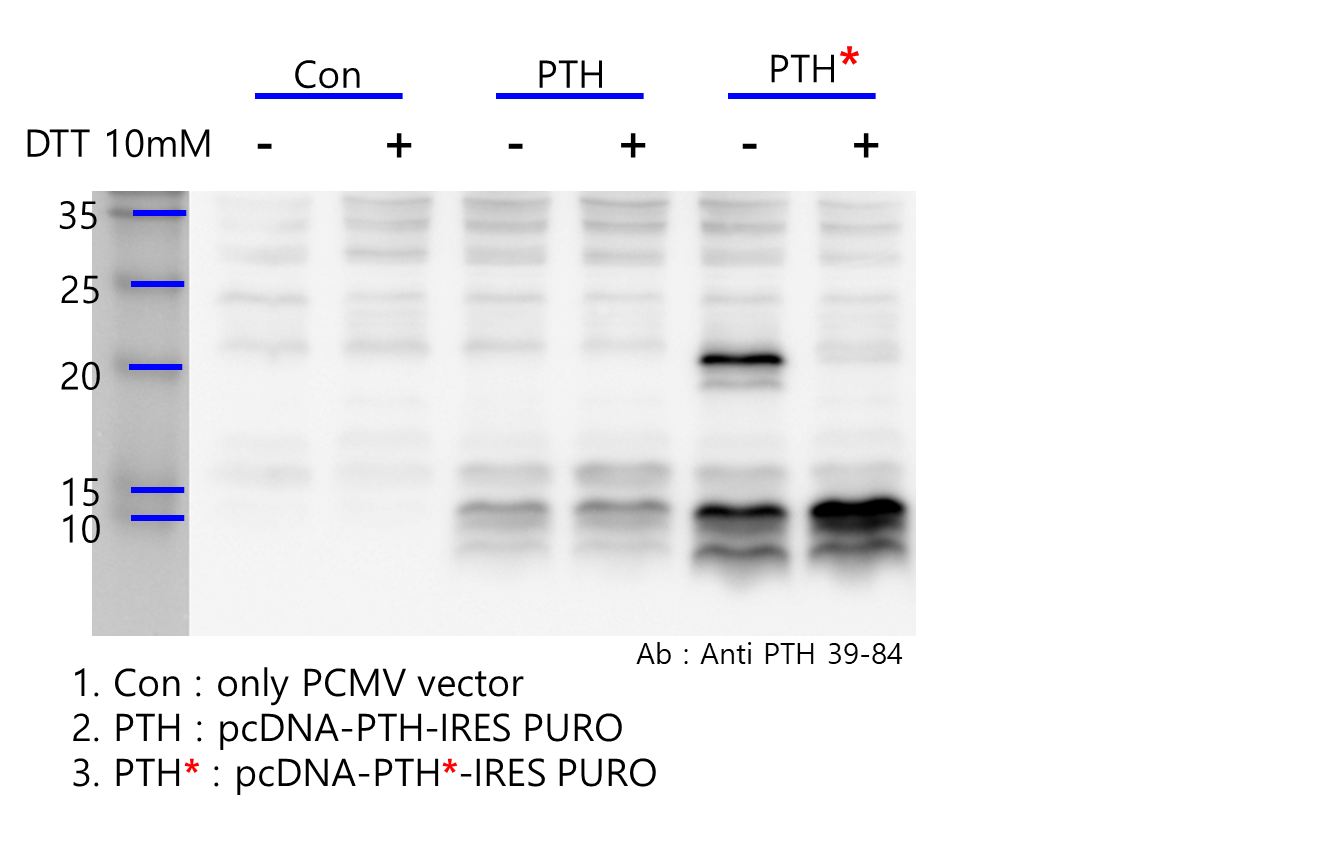

**Figure supplement 2.** Identification of dimeric ^R25C^PTH (1-84) peptide.

HEK293T cells were separately transfected with three plasmids, pcDNA3.0 empty vector, pcDNA3.0-(pre-pro-PTH)-IRES, and pcDNA3.0(^R56C^pre-pro-PTH)-IRES. Total cell lysates were extracted from each transfected group and divided into two types of samples, reduced and non-reduced. The results revealed a specific dimeric formation of ^R25C^PTH(1-84) (QuidelOrtho, 21-3010, PTH Antibody (Center 39-84)). The primary antibody used in this experiment binds to the 39-84 region of mature PTH(1-84). 10 mM DTT was used for reducing agent.
