## Supplementary material for "Dimeric ^R25C^PTH(1-34) Activates the Parathyroid Hormone-1 Receptor *in vitro* and Stimulates Bone Formation in Osteoporotic Female Mice": Figure supplement 3

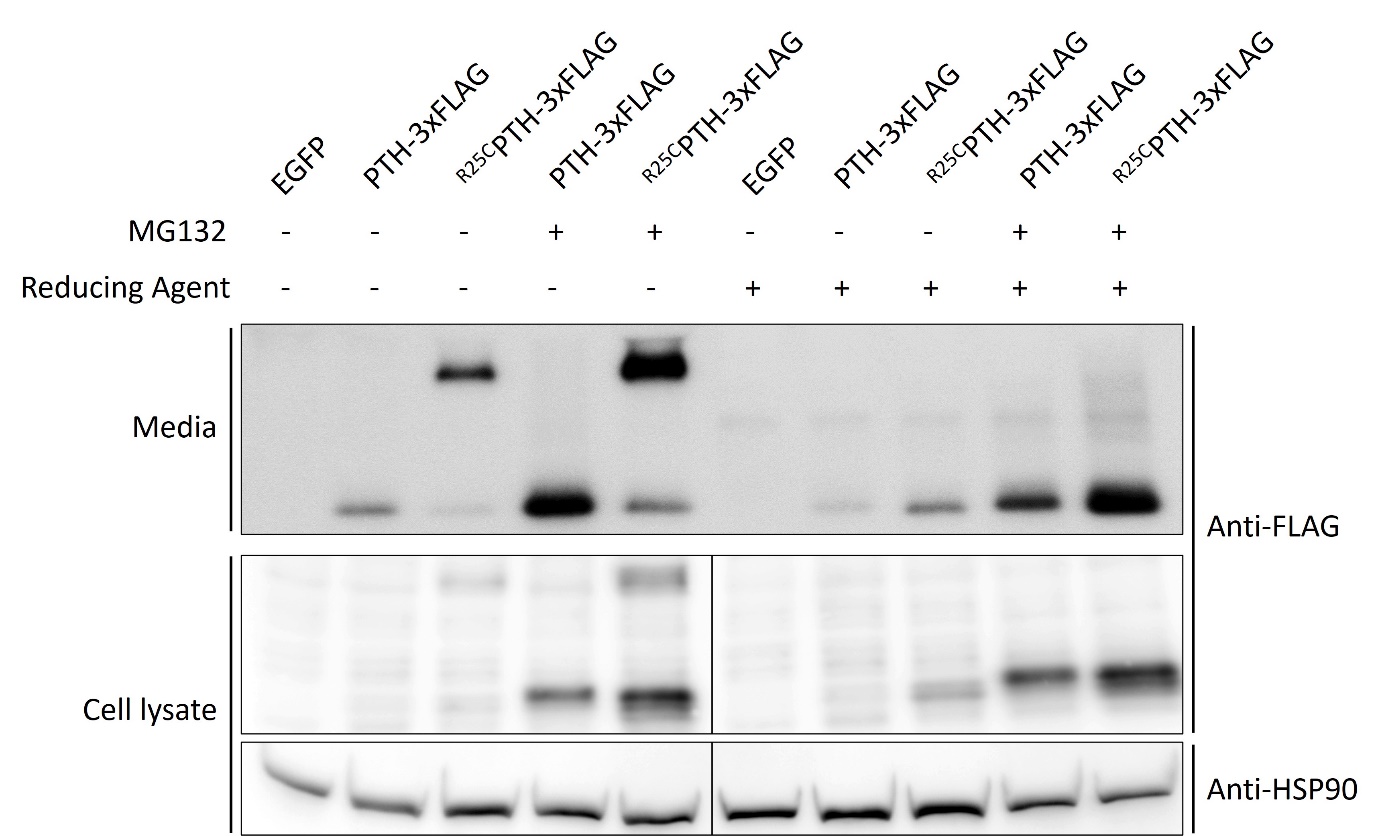

**Figure supplement 3.** Influence of proteasome inhibitor MG132 on PTH and ^R25C^PTH stability

To validate whether inhibition of proteasome-mediated degradation can restore the decreased expression level of wild-type PTH compared to ^R25C^PTH under normal conditions, a proteasome inhibition assay using MG132 was conducted. HEK293T cells were transfected with plasmids including control vector, pcDNA3.1-PTH-3xFLAG, and pcDNA3.1-^R25C^PTH-3xFLAG. After 24 hours of transfection, MG132 was administered for 24 hours, and media as well as total cell lysate were collected for subsequent western blot analysis. Each sample was divided into two types, reduced and non-reduced. The results indicated that both PTH and ^R25C^PTH exhibited increased protein expression levels following MG132 treatment-induced proteasome inhibition. However, the disparity in expression levels between the PTH and ^R25C^PTH remained unchanged.
