## Supplementary Table 1 for "Dimeric ^R25C^PTH(1-34) Activates the Parathyroid Hormone-1 Receptor *in vitro* and Stimulates Bone Formation in Osteoporotic Female Mice"

Supplementary Table 1. List of primers used in this study

| Name | Sequence | Description |
| --- | --- | --- |
| PTH-Forward | 5’-GGGGACAACTTTGTACAAAAAAGTTGGCATGATACCTGCAAA  AGACATGGCTAAAG-3’ | Plasmid construction |
| PTH-Reverse | 5’-GGGGACAACTTTGTACAAGAAAGTTGGGTACTGGGATTTAGC  TTTAGTTAATACATTCACATCAG-3’ | Plasmid construction |
